# Genome related features of introns and exons reveal new properties and pareto fronts

**DOI:** 10.1101/2023.09.20.558659

**Authors:** Yekbun Adiguzel

**Affiliations:** Atilim University

**Keywords:** Genome annotation, length feature, count feature, eukaryotes

## Abstract

Genome annotations reveal vast amount information, as well as genome related features. We looked at the genome related features of eukaryotes, grouped as primates, rodents, birds, fishes, insects, and plants. Features related to introns and exons reveal distinct assets. Genes’ mean lengths are positively correlated with the collective data of counts of exons in coding transcripts and both the mean lengths and counts of introns in coding transcripts. Mean lengths of introns in non-coding transcripts are also positively correlated and correlations with the counts-data are excluding the data of plants and fishes. These are valid for the median lengths as well. In addition, certain feature comparisons reveal separation of primates, rodents, birds, fishes, insects, and plants. We also observed pareto fronts in the cumulative data with the max length values of genes and introns in coding transcripts, as well as introns in coding and non-coding transcripts.

## Introduction

Shortly before this century, the first genome sequence belonging to the unicellular organism *Haemophilus influenzae* was published (Devos and Valencia, 2001; Armengaud, 2009). At the end of 2000, 3 eukaryote genomes were sequenced (Bernal, et al., 2001; Aubourg and Rouzé, 2001), and more were in progress (Rust, et al., 2002). In 2009, 65 eukaryote genomes were established (Armengaud, 2009), while at the end of 2017, up to 4963 eukaryote genomes were available at NCBI (Liu, et al., 2018). In 2019, 8987, and in 2023, 30530 eukaryote genomes were available at NCBI, revealing an exponential increase (https://www.ncbi.nlm.nih.gov/genome/browse#!/overview/; accessed on August 20, 2019, and September 20, 2023). Meaningful information (Abril and Castellano Hereza, 2019) can be extracted from great number of sequenced-data by predictions (Aubourg and Rouzé, 2001; Koonin and Galperin, 2003). Distant evolutionary relationships are inferred from structural similarities (Furnham, et al., 2012). Here, we derive exons’, and introns’ related information from the feature lengths of the annotation reports of primates, rodents, birds, fishes, insects, and plants, which are all among the eukaryotes.

Exon-intron structure of eukaryotic genes has long been a popular evolutionary biology matter. About up to 95% of the mammalian primary protein coding transcripts’ length is introns (Mattick and Gagen, 2001). Many ancestral introns are persisting since the last eukaryotic common ancestor, but unicellular eukaryotes are intron-poor while multicellular eukaryotes are intron-rich (Rogozin, et al., 2012). In this study, we are dealing with features like the sizes and counts. Intron counts is related to the intron dynamics in the eukaryote evolution where the last common ancestor of eukaryotes contained more than 2.15 introns per kilobase and the last common ancestor of multicellular lifeforms contained about 3.4 introns per kilobase, followed by a steeper decline in the rate of gain than the loss of introns, in the last ∼1.3 billon years (Carmel, et al., 2007). Another study (Irimia, et al., 2007) also indicated that the ancestral eukaryotes had relatively large intron numbers and there is association between the splice site sequences and the spliceosomal intron numbers such that the sequences are strongly conserved when the intron number is low. They indicated additionally that the ancestral eukaryotes had weak splice sites. In support, another work revealed that weak splice sites are keeping introns short while “strong splice sites allow recognition of exons flanked by long introns” and this phenomenon appeared during vertebrate evolution (Gelfman, et al., 2012). Alternative splicing is a phenomenon that leads to the production of different protein isoforms (Wang, et al., 2008). There are even conserved alternative exons indictive of tissue-specific, conserved splicing regulators, in a subset of tissues, and lineage specific manner (Merkin, et al., 2012). Alternatively spliced novel exons are tended to be in introns longer than 1000 nucleotides (Roy, et al., 2008). This large intron sizes are relatively few in the plant introns, which are expansion-limited, and they are transposon-free, contrary to those in animals (Wu, et al., 2013). In a review, long introns are suggested ((Comeron, et al., 2008) cited in (Jo and Choi, 2015)) to be favored by increasing the efficiency of natural selection through decreasing the Hill-Robertson interference, which is the “genetic linkage between two sites under selection in finite populations.” Other studies (Gorlova, et al., 2014) also reported similar findings. On the other hand, introns are inherently related to the transcript lengths and natural selection suppresses changes for genes with longer transcripts (Lopes, et al., 2021). Longer transcripts is also the case for the transcripts that are expressed at the early developmental stages rather than those expressed as fast responses (Lopes, et al., 2021). Another intriguing finding is that the intron lengths in genes associated with developmental patterning are conserved (Keane and Seoighe, 2016). In one of our earlier studies, we attempted to relate the lengths of encoding DNA with the encoded proteins through the introns, with a theoretical model (Adiguzel, 2022).

Researchers delineated the factors influencing the intron size as the transposons, sizes and frequencies of deletion events, and factors controlling gene expression and regulation like the presence of regulatory elements and RNA genes, and alternative splicing (Zhu, et al., 2009), based on earlier studies (Duret, 2001; Bartolome, et al., 2002; Petrov, et al., 2000; Maxwell and Fournier, 1995; Xing and Lee, 2006; Bergman and Kreitman, 2001). Besides, these researchers investigated the lengths, divergences, and GC contents of introns and exons in an ordinal position dependent manner. They found that the ordinal position of introns are exons are inversely related to their corresponding lengths and divergences, as well as co-variation of the investigated parameters in the flanking introns and exons. Also, average exon (or intron) length, GC content, and divergence decreased with the total number of exons. GC contents of introns were more complex.

Intriguingly, introns to intergenic sequence ratios were found to be similar across animal genomes while that of the model organisms were close to 1, which was interpreted as a possible under annotation of the non-model organisms (Francis and Wörheide, 2017). In relation, absolute lower or upper limits, which will be presented also in this study, are inherently related to the observation of the relation of genome sizes with the cell sizes, where there is an absolute lower limit of the genome size (Malerba, et al., 2020). Regarding the relation of introns with genome sizes, Lozada-Chávez and coworkers (Lozada-Chávez, et al., 2018) argued that sizes and repeat compositions of introns are only weakly scaling with genome size changes at the broadest phylogenetic scale. Those phylogenetic scales ranging between fungi, plants, stramenophiles, holozoans, chlorophytes, alveolates, and amoebozoans, are not readily comparable to our study, which is presented below.

## Results and discussion

### A. Genes’ median and mean lengths vs exons’ and introns’ parameters

#### 1. Not exons’ but introns’ mean lengths increase with the genes’ mean lengths

##### a. Exons’ mean lengths do not increase with the genes’ mean lengths

Genes’ mean lengths *vs* mean lengths of exons in coding (Fig. 1b) and non-coding (Fig. 1c) transcripts reveal that the mean lengths of exons in coding transcripts are constant with similar variations. Corresponding boxplots of the mean lengths of exons in coding transcripts are displayed in Fig. 1f. The data ranges are close, within 100–600 nucleotides (nt) and populated at about 100–400 nt (Fig. 1f).

**Fig. 1.**
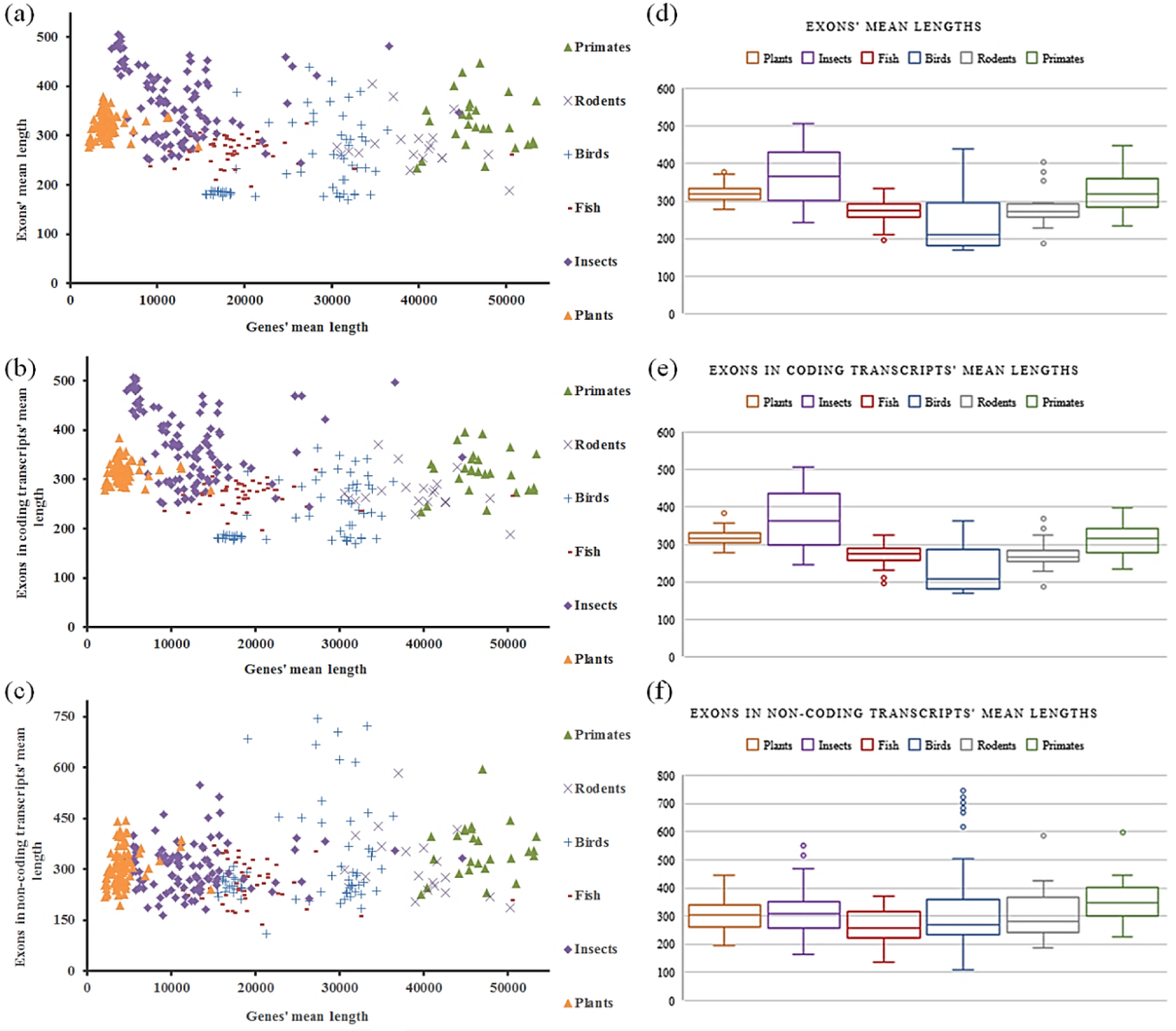
Scatter plots of the mean lengths of genes *vs* exons (a), genes *vs* exons in coding transcripts (b), genes *vs* exons in non-coding transcripts (c); and boxplots of the mean lengths of exons (d), exons in coding transcripts (e), and exons in non-coding transcripts (f), in eukaryotes.

Exons’, exons in coding transcripts’, and exons in non-coding transcripts’ mean lengths of separate groups are displayed in Table 1. Accordingly, exons’ mean lengths of:

**Table 1.**
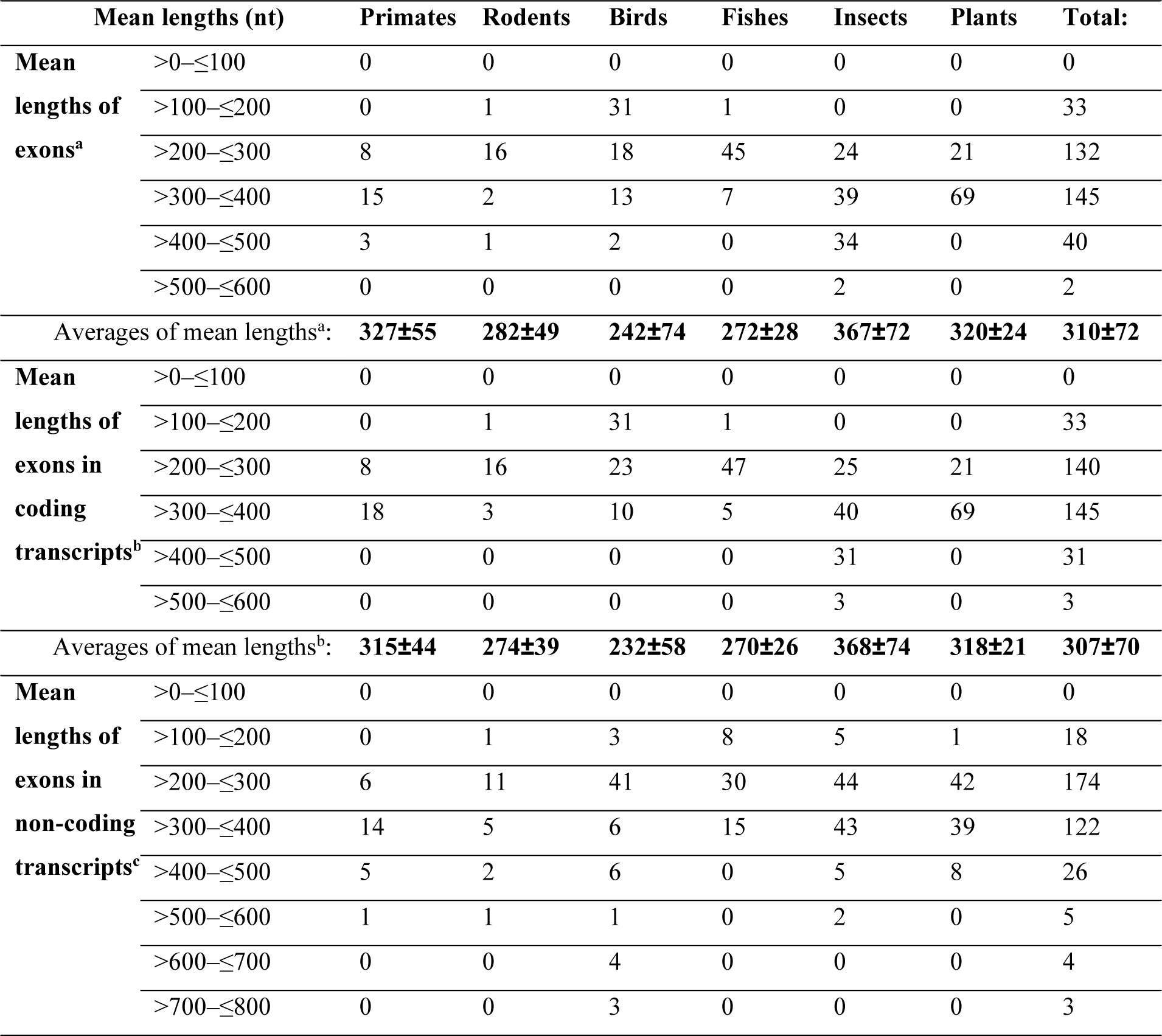
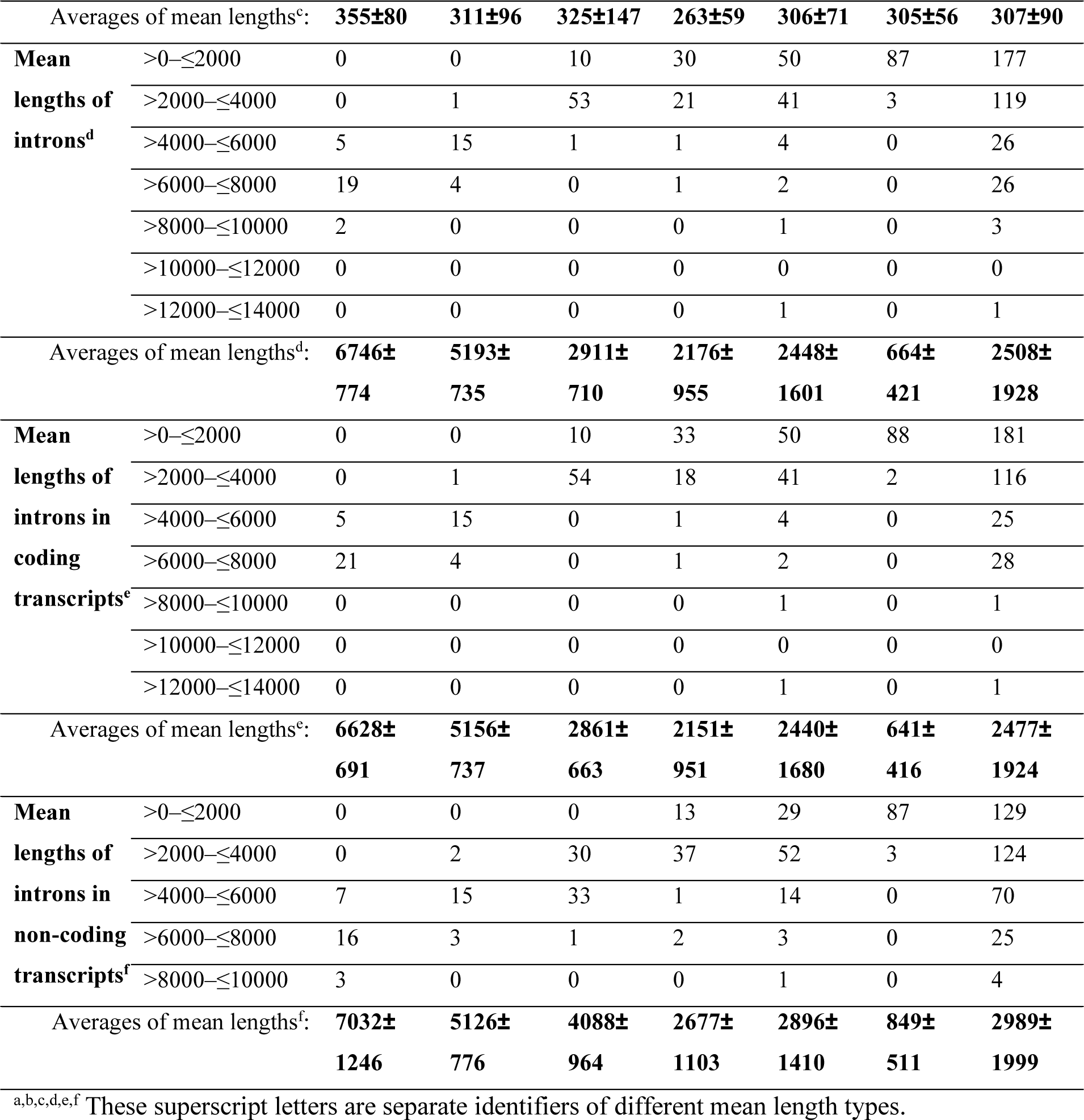
The numbers of individual species in different ranges of mean lengths of exons, introns, and those in the coding and non-coding transcripts. Calculated-averages of each group is written in bold and at the end of the corresponding data-group.

Primates are within the 200–500 nt range, and 15/26 are 300–400 nt,

Rodents are within the 100–500 nt range, and 16/20 are 200–300 nt,

Birds are within the 100–500 nt range, and 31/64 are 100–200 nt,

Fishes are within the 100–400 nt range, and 45/53 are 200–300 nt,

Insects are within the 200–600 nt range, and 39/99 are 300–400 nt,

Plants are within the 200–400 nt range, and 69/90 are 300–400 nt.

Average of the exons’ mean lengths is 310±72 nt while that of exons in coding transcripts is 307±70 nucleotides, and of exons in non-coding transcripts is 307±90 nt (Table 1).

The mean lengths of exons (Fig. 1a,d) are like that in coding transcripts (Fig. 1b,e) while that in non-coding transcripts are similar in values, but diverging more (Fig. 1c,f). This indicates the abundance of exons in the coding transcripts, which is observed in the respective data of counts only, in a comparative manner (Fig. 2a,b). Counts of exons in coding transcripts are close to that of exons and positively correlated (Fig. 2a) while counts of exons in non-coding transcripts are about 5 times less and barely correlated as a holistic data (Fig. 2b). Therefore, we rather mention only the features of plots with exons in coding transcripts rather than both exons and exons in coding transcripts. The situation is the same for introns (Fig. 2c,d).

**Fig. 2.**
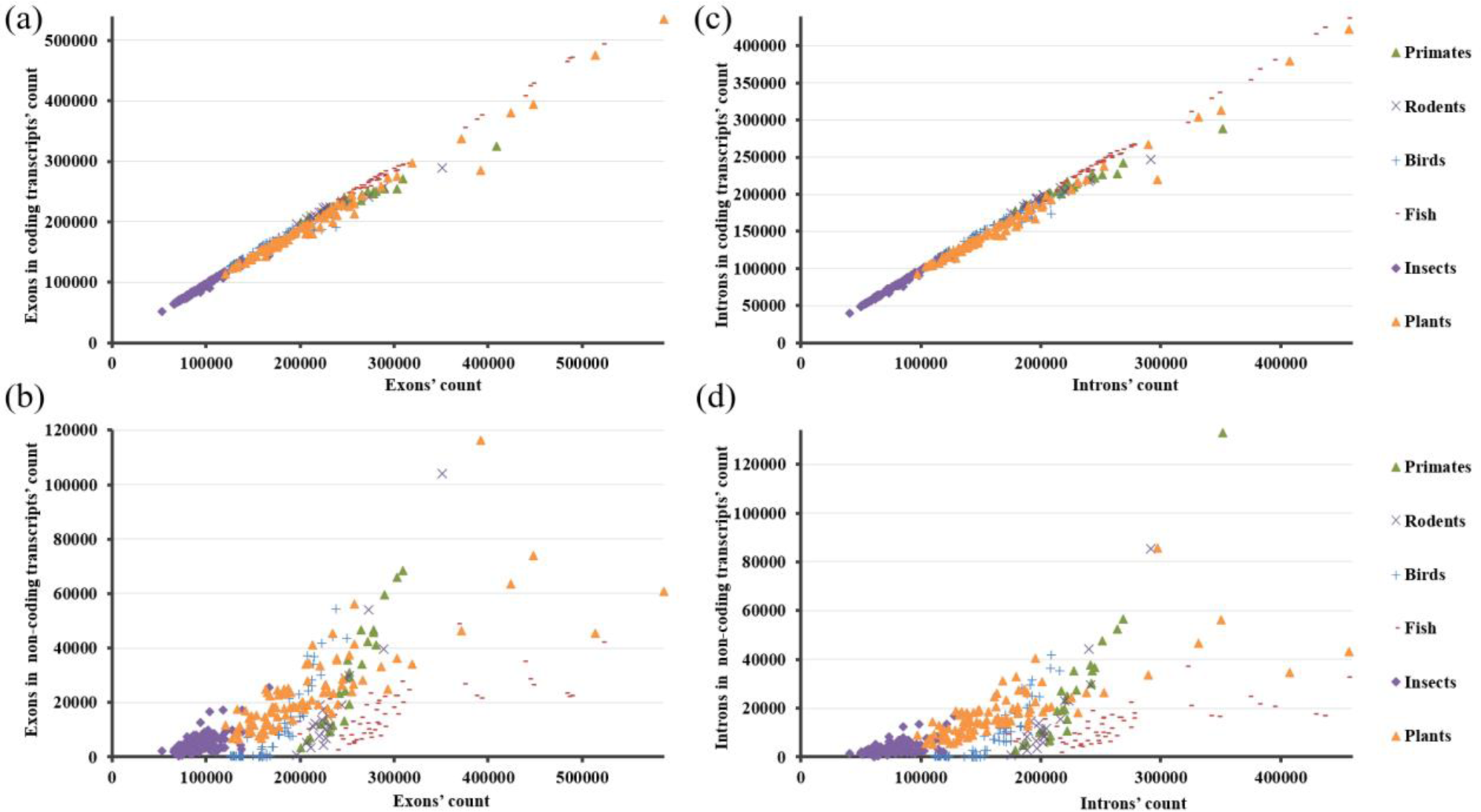
Scatter plots of the counts of exons *vs* exons in coding transcripts (a), exons *vs* exons in non-coding transcripts (b), introns *vs* introns in coding transcripts (c), and introns *vs* introns in non-coding transcripts (d), in eukaryotes.

Linear regression analysis of the data in Fig. 1a, genes’ mean lengths *vs* exons’ mean lengths, reveals that only the slope of the insects’ data is significantly non-zero (supplementary information (Supplemental_Table_S1). That is the case also for the data in Fig. 1b, i.e., genes’ mean lengths *vs* exons in coding transcripts’ mean lengths (Supplemental_Table_S2). Exons are dominated by those in the coding transcripts, like in case of introns. Accordingly, these results and the other results revealing similarity of the dependences of the exons or introns and those in the coding transcripts, correspondingly, are presented in the figures as the proof-of-the-concept. However, as mentioned above, they are not separately discussed.

In case of the genes’ mean lengths *vs* exons in non-coding transcripts’ mean lengths (Fig. 1c), even the slope of the data of insects is not significantly non-zero (Supplemental_Table_S3). Overall, mean lengths of exons, and that of the related parameters, are not revealing any correlated change with the mean lengths of genes. In contrast, each of the mean lengths of introns and the mean lengths of introns’ related parameters are positively correlated with the genes’ mean lengths (Fig. 3). Linear regression analysis of the genes’ mean lengths *vs* introns’ mean lengths of the cumulative data of all groups, reveals that the slope is significantly non-zero with 0.83 goodness of fit (Supplemental_Table_S4). That is the case also for the genes’ mean lengths *vs* introns in coding transcripts’ mean lengths and genes’ mean lengths *vs* introns in non-coding transcripts’ mean lengths, wherein the slopes are again significantly non-zero, with 0.81 and 0.77 goodness of fits, respectively (Supplemental_Table_S5 and Supplemental_Table_S6).

**Fig. 3.**
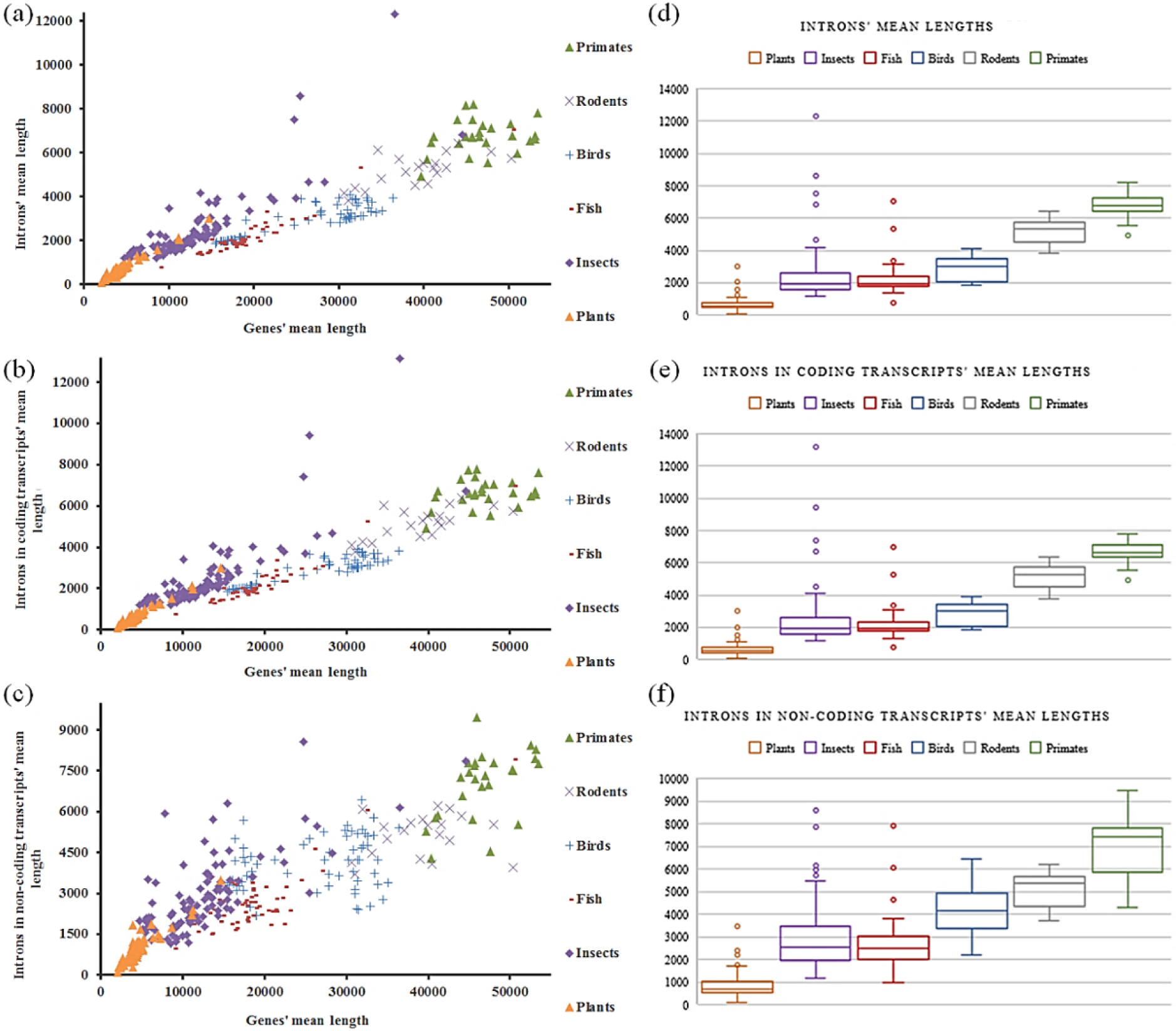
Scatter plots of the mean lengths of genes *vs* introns (a), genes *vs* introns in coding transcripts (b), genes *vs* introns in non-coding transcripts (c); and boxplots of the mean lengths of introns (d), introns in coding transcripts (e), and introns in non-coding transcripts (f), in eukaryotes.

##### b. Introns’ mean lengths increase with the genes’ mean lengths

Introns-related data is observed to be varying up to a mean length of 8000 nt, although there are extremes with mean lengths well above that (Table I and Fig. 3). Averages of the mean lengths of introns (2508±1928 nt) and introns in coding transcripts (2477±1924 nt) are about 8 times more than that of the exons and exons in coding transcripts, and that of the introns in non-coding transcripts (2989±1999 nt) is about 10 times more than that of the exons in non-coding transcripts. Except for the rodents’ data, averages of the mean lengths of introns in non-coding transcripts are higher than that of the introns and introns in coding transcripts when the respective data-groups are compared (Table I). Mean lengths of introns (Fig. 3a,d) are like the mean lengths of introns in coding transcripts (Fig. 3b,e) while that of the non-coding transcripts are similar in values but diverging more (Fig. 3c,f). This similarity indicates abundance of introns in coding transcripts, which is observed in the counts (Fig. 4d-f).

**Fig. 4.**
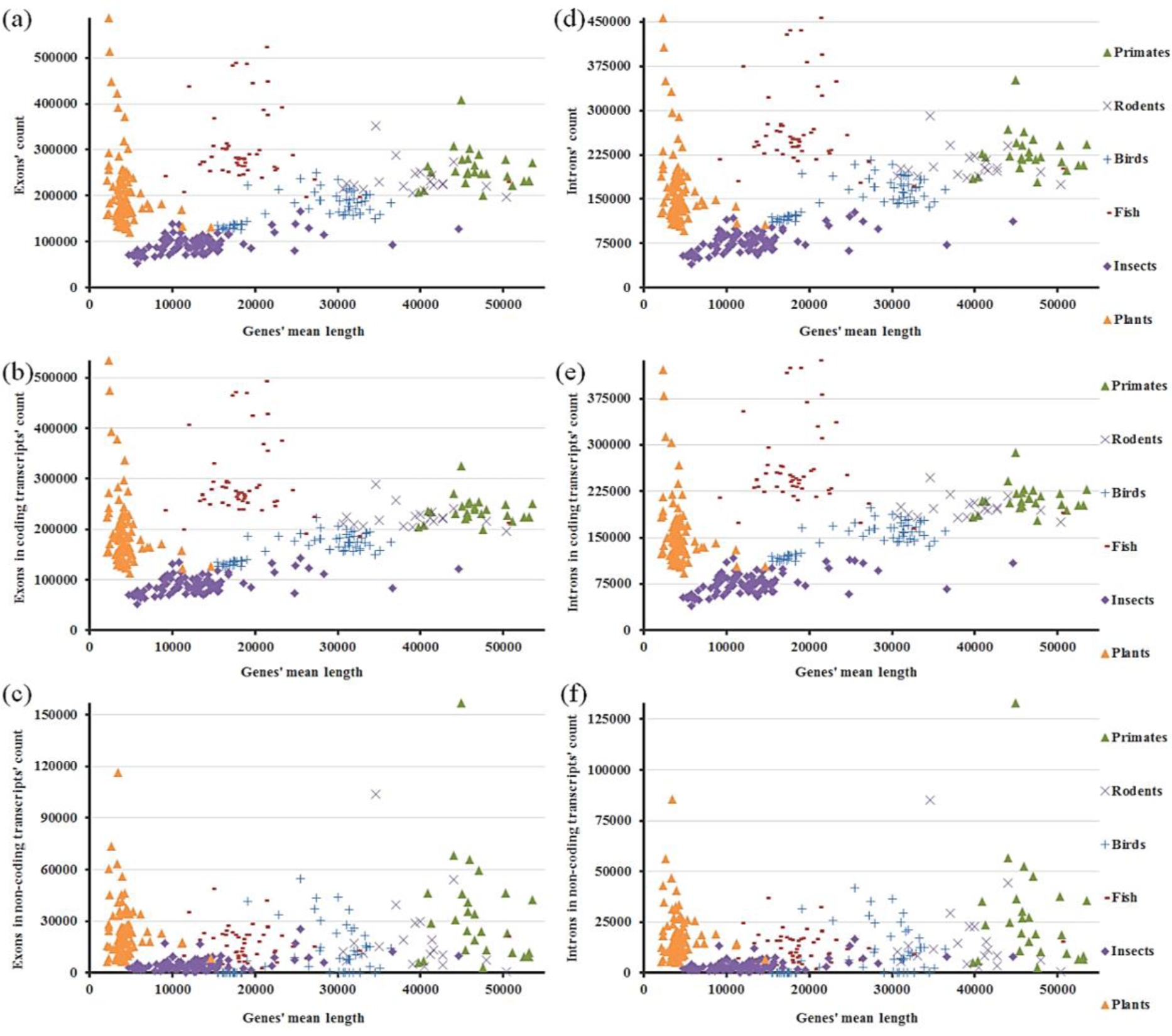
Scatter plots of the mean lengths of genes *vs* counts of exons (a), mean lengths of genes *vs* counts of exons in coding transcripts (b), and mean lengths of genes *vs* counts of exons in non-coding transcripts (c), mean lengths of genes *vs* counts of introns (d), mean lengths of genes *vs* counts of introns in coding transcripts (e), mean lengths of genes *vs* counts of introns in non-coding transcripts (f), in eukaryotes.

As can be observed in Table I and Fig. 3, there is inter-group variation in the data of introns, and the related parameters. There is also positive correlation of the mean lengths of introns and the related parameters with the genes mean lengths (Fig. 3) at the levels of both the inter-species’ groups, and the intra-species’ groups.

#### 2. Counts of exons and introns in coding transcripts increase with the genes’ mean lengths

Order of the groups with increasing counts in association with the genes’ mean lengths are as follows: insects, birds, rodents, and primates (Fig. 4). As mentioned, mean lengths of introns and the related parameters are also positively correlated (Fig. 3), and the corresponding order together with the data of plants and fishes is as follows: plants, insects, fishes, birds, rodents, and primates (Fig. 3).

Except for the plants and fishes, positive correlation with the genes’ mean lengths is present for the counts of exons and introns, which are in coding transcripts (Fig. 4b,e), but absent for the exons in non-coding transcripts and introns in non-coding transcripts (Fig. 4c,f). Linear regression analyses of the combined data except that of plants and fishes reveal that the slopes of the genes’ mean lengths *vs* counts-data of either exons or exons in coding transcripts are significantly non-zero with 0.77 and 0.82 goodness of fits, respectively (Supplemental_Table_S7 and Supplemental_Table_S8). The same analyses for the genes’ mean lengths *vs* counts-data of either introns or introns in coding transcripts also reveal that the slopes are significantly non-zero with 0.79 and 0.83 goodness of fits, respectively (Supplemental_Table_S9 and Supplemental_Table_S10). That correlation is visibly lost in the genes’ mean lengths *vs* counts of exons and introns, which are in non-coding transcripts (Fig. 4c,f). Linear regression analyses of the combined data except that of plants reveal that the slopes of the genes’ mean lengths *vs* counts-data of either exons in non-coding transcripts or introns in non-coding transcripts are still significantly non-zero, but with a low (0.19) goodness of fit (Supplemental_Table_S11 and Supplemental_Table_S12).

In sum, when plotted against the genes’ mean lengths, mean lengths of exons in coding transcripts are relatively constant (Fig. 1b) while their counts are correlated with the genes’ mean lengths except for that of plants and fishes (Fig. 4b). However, exons in non-coding transcripts’ mean lengths are still relatively constant with respect to the genes’ mean lengths (Fig. 1c) and this time, the situation complies with that of the counts of exons in non-coding transcripts except for that of plants (Fig. 4c). Mean lengths of introns in coding transcripts are correlated with the genes’ mean lengths (Fig. 3b), like their counts’ relation with the genes’ mean lengths except for that of plants and fishes (Fig. 4e). However, introns in non-coding transcripts’ mean lengths are also correlated with the genes’ mean lengths (Fig. 3c) while their counts appear constant except for that of plants (Fig. 4f).

These findings suggest that mean lengths of exons in coding transcripts are rather constant when they are plotted against the genes’ mean lengths (Fig. 1b,e). However, counts of exons in coding transcripts are positively correlated with the genes’ mean lengths, except for the plants and fishes (Fig. 4b). Exons in non-coding transcripts’ mean lengths are also relatively constant regarding the genes’ mean lengths (Fig. 1c,f). Yet, genes’ mean lengths *vs* exons in non-coding transcripts’ counts also appear constant (Fig. 4c). Differently, genes’ mean lengths *vs* mean lengths of introns in coding transcripts are positively correlated (Fig. 3b,e) and there is positive correlation between the genes’ mean lengths and the counts of introns in coding transcripts, except for that of plants and fishes (Fig. 4e). Introns in non-coding transcripts’ mean lengths are also correlated with the genes’ mean lengths (Fig. 3c,f), but the respective counts do not reveal such a relation with the genes’ mean lengths (Fig. 4f).

As stated earlier, we mention only those in the coding and non-coding transcripts instead of mentioning additionally the introns and exons that are contained in both the coding and the non-coding transcripts but dominated by those in the coding transcripts. Counts of exons in coding transcripts except for that of plants and fishes are positively correlated with the genes’ mean lengths (Fig. 4b). Both mean lengths (Fig. 3b,e) and counts of introns in coding transcripts are positively correlated (Fig. 4e). Mean lengths of introns in non-coding transcripts are also positively correlated with the genes’ mean lengths (Fig. 3c,f). Correlations with the counts of introns-related data are also excluding the plants’, and fishes’ data.

Plots of genes’ *vs* exons-parameters’ mean lengths are similar (Fig. 1). This is the case for the comparable plots with the introns’ data (Fig. 3), but plots in Fig.1 and Fig. 3 are divergent. However, exons- and introns-related comparable plots with the counts-data are similar (Fig. 4). Namely, in relation with the genes’ mean lengths, mean lengths of introns and exons behave differently, but not their counts. Also of note, lengths of the introns are more, in general, and exons or introns in coding transcripts are abundant when counts in coding and non-coding transcripts are compared.

The discussion up to here can be summarized as follows: Counts of exons in coding transcripts, rather than their mean lengths, are correlated with the genes’ mean lengths. Introns in coding transcripts’, both mean lengths and counts, and introns in non-coding transcripts mean lengths, are correlated with the genes’ mean lengths. As mentioned, correlations with the counts exclude fishes and plants. Present observation is indicative of a size constriction in terms of the mean lengths of exons both in coding and non-coding transcripts. There is no such limitation in case of the introns-related parameters. Excluding the fishes’ and plants’ data, counts of both exons and introns, both in the coding transcripts, are positively correlated with the genes’ mean lengths. In other words, *exons mean lengths are limited* and independent of the genes’ mean lengths but counts of exons in coding transcripts increase with that. Introns in coding transcripts both mean lengths and counts, and introns in non-coding transcripts’ only mean lengths, increase with the genes’ mean lengths. In this sense, there is a *limitation of the counts of exons and introns in non-coding transcripts*. There, plants’ data is following the same distinctive pattern when correlation was observed while the separation of fishes’ data from the others ceases.

Genes’ mean lengths are calculated as well, as described in the methods section. As a result, only 7 species among 352 revealed to have less than 70% of the lengths of the specified mean lengths of genes within the annotations. Among those 7 species, calculation results of 6 of those are having 60–70% of the lengths of the specified mean lengths of genes while 1 is having 50–60% of the length of the specified mean length of genes.

One can see roughly the higher contributing component in the genes, by subtracting the product of introns’ mean lengths and their counts per gene, from the product of the exons’ mean lengths and their counts per gene. Accordingly, only16 species revealed to have a positive result, suggesting that the total lengths of introns are higher in general. When we take the ratio of the calculated total intron-lengths per gene and the calculated mean length of genes, the average of the results is 0.80±0.15. So, the introns’ lengths are about 80% of the total lengths of the genes, in average. However, counts of exons are higher in number than the counts of introns. This is supporting higher average lengths of introns, leading to a higher total length of introns per gene despite a higher number of exons per gene. Increase in the ratio of the total length of introns per gene length is not linear, but logarithmic, and approaching 1 (Fig. 5). Specifically, as mean sizes of genes increase, ratios of the total lengths of introns per gene lengths initially increase fast, which is in case of plants with the lowest mean lengths of genes, and then the ratios increase gradually slower and approaches to 1, in the order of insects, fishes, birds, rodents, and primates. This type of increase is due to the simultaneous increases in the mean lengths of the genes and the counts of introns and exons, both in the coding transcripts, and the lengths of the introns. Same rates of increases in the exons in the coding transcripts and introns in the coding transcripts (Fig. 4) results in a constant ratio of the lengths of introns and exons, considering constant lengths.

**Fig. 5.**
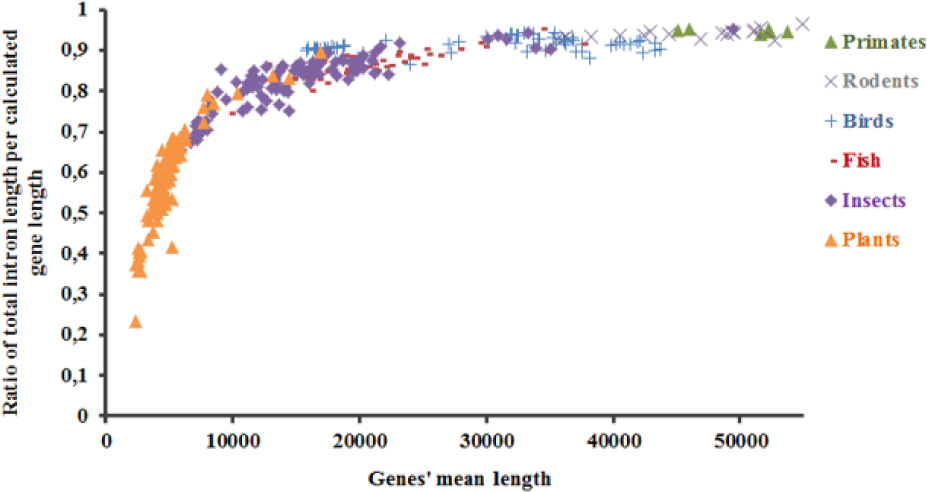
Scatter plot of the mean lengths of genes *vs* total lengths of the introns per genes lengths, in eukaryotes. Calculated values of the mean lengths of genes are used. Calculations are performed with the data from (Adiguzel, 2023).

### Additional observations

Corresponding plots with median lengths reveal similarities to those discussed until here (Fig. 6). However, negative correlation of the exons in coding transcripts’ median lengths with the genes’ mean length is well observed (Fig. 6b), which was not readily observed in the corresponding plots with the mean lengths (Fig. 1b). Besides, especially the positive correlation of the median lengths of genes and that of introns in coding transcripts (Fig. 6d,e) reveal a phenomenon termed as Simpson’s paradox. Simpson’s paradox (Simpson, 1951) is “a statistical phenomenon in which an observed association between two variables at the population level (e.g., positive, negative, or independent) can surprisingly change, disappear, or reverse when one examines the data further at the level of subpopulations” (Bonovas and Piovani, 2023). Regarding Simpson’s paradox, intra-group variations follow a distinct pattern than that of the combined data in Fig. 6d,e. Namely, median lengths of the genes and that of introns in coding transcripts, except for the plants and insects, are having a slight positive correlation while the positive correlation of the combined data is stronger (Fig. 6d,e). This observation is less evident in the median lengths of genes *vs* that of introns in non-coding transcripts (Fig. 6f). If we also compare the plots with the counts data as well, corresponding plots of that displayed in Fig. 4 is similar when the plots contain median lengths of genes instead of the mean lengths (Supplemental_Fig_S1, page 18, first row). Also, distinct patterns of the intra-group variations from the combined data is the case in the other plots as well, e.g., in the median length plots of exons in coding transcripts *vs* exons in non-coding transcripts (Supplemental_Fig_S1, page 3).

**Fig. 6.**
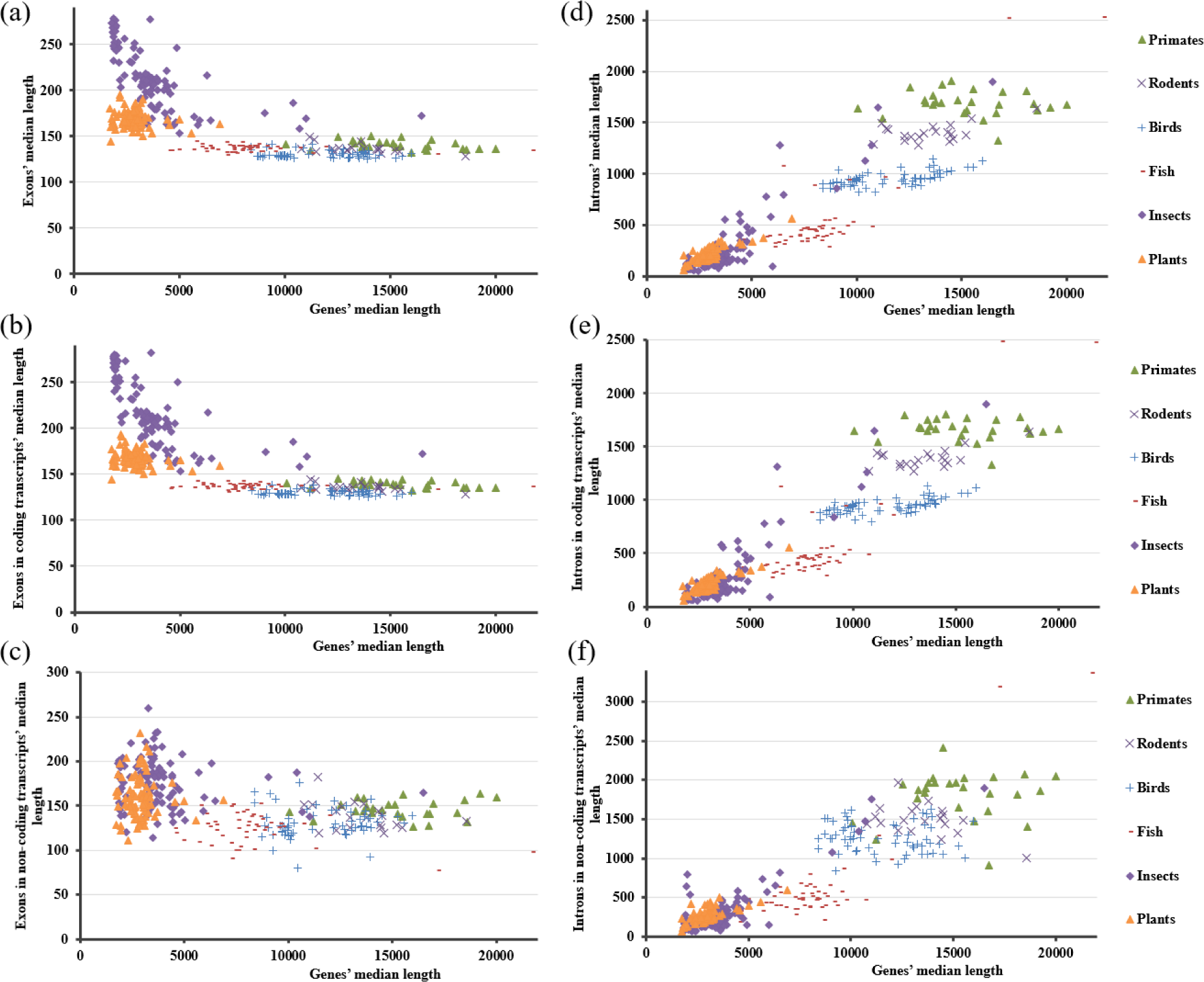
Scatter plots of the median lengths of genes *vs* exons (a), genes *vs* exons in coding transcripts (b), genes *vs* exons in non-coding transcripts (c); genes *vs* introns (d), genes *vs* introns in coding transcripts (e), genes *vs* introns in non-coding transcripts (f), in eukaryotes.

At the genomic and proteomic levels, current findings also suggest that the introns’ mean lengths can be related with that of the expressed proteins, through the sizes of the mature mRNAs, and indeed through the total mean lengths of exons per gene. This is based on the observation that exons in coding transcripts’ only counts, but not the lengths, increase with the genes’ mean lengths. This would increase the introns in coding transcripts’ counts as well because exons are connected to each other with introns. However, increase in the mean lengths of both introns in coding transcripts and introns in non-coding transcripts along with the genes’ mean lengths suggest positive correlation of the introns’ mean lengths with the protein sizes. This observation is inherent to the relevant assumption underlying in one of our earlier studies (Adiguzel, 2022), namely that assumption is justified here. Knowing that the number of introns, exons, and intron lengths in genes are variable. This correlation suggested by the results is valid for the genomes and proteomes of the individual species.

As mentioned at the introduction, longer transcripts is the case for the transcripts that are expressed at the early developmental stages rather than those expressed as fast responses (Lopes, et al., 2021). Namely, there is transcript length variation along with the tissue expression sites and the expression times. In this sense, widening of the transcript length scales, which is related to the intron-parameters in turn, can be indicative of the scope of variation in the tissue-dependent expressions and the responses.

### B. Gorup separation with exons’ and introns’ parameters

When the introns-, exons-related data are utilized, certain plots of the mean or median lengths *vs* counts reveal separation of groups as primates, rodents, birds, fishes, insects, and plants (Fig. 7). There are only singular data entering the domains of the other groups, yet the separations are successful, as observed in Fig. 7. Accordingly, Fig. 7a,b display counts of exons *vs* median lengths of introns, and introns in coding transcripts, respectively. Fig. 7c is the counts of exons in coding transcripts *vs* median lengths of introns in coding transcripts. Figures on their right (i.e., Fig. 7f,g,h) are the corresponding plots with the mean values instead of the medians. Fig. 7d,e are the counts of introns in coding transcripts plotted against the mean and median lengths of introns, respectively, while Fig. 7i,j are the counts of introns *vs* mean and median lengths of introns in coding transcripts, respectively. Accordingly, figures at the last two rows (Fig. 7d,e,i,j) present combinations of the plots of the mean and median lengths, and counts, of introns and introns in coding transcripts. Remaining figures at the top three rows (Fig. 7a,b,c,f,g,h) involve only counts as the exons-related feature, but not any length-feature of exons.

**Fig. 7.**
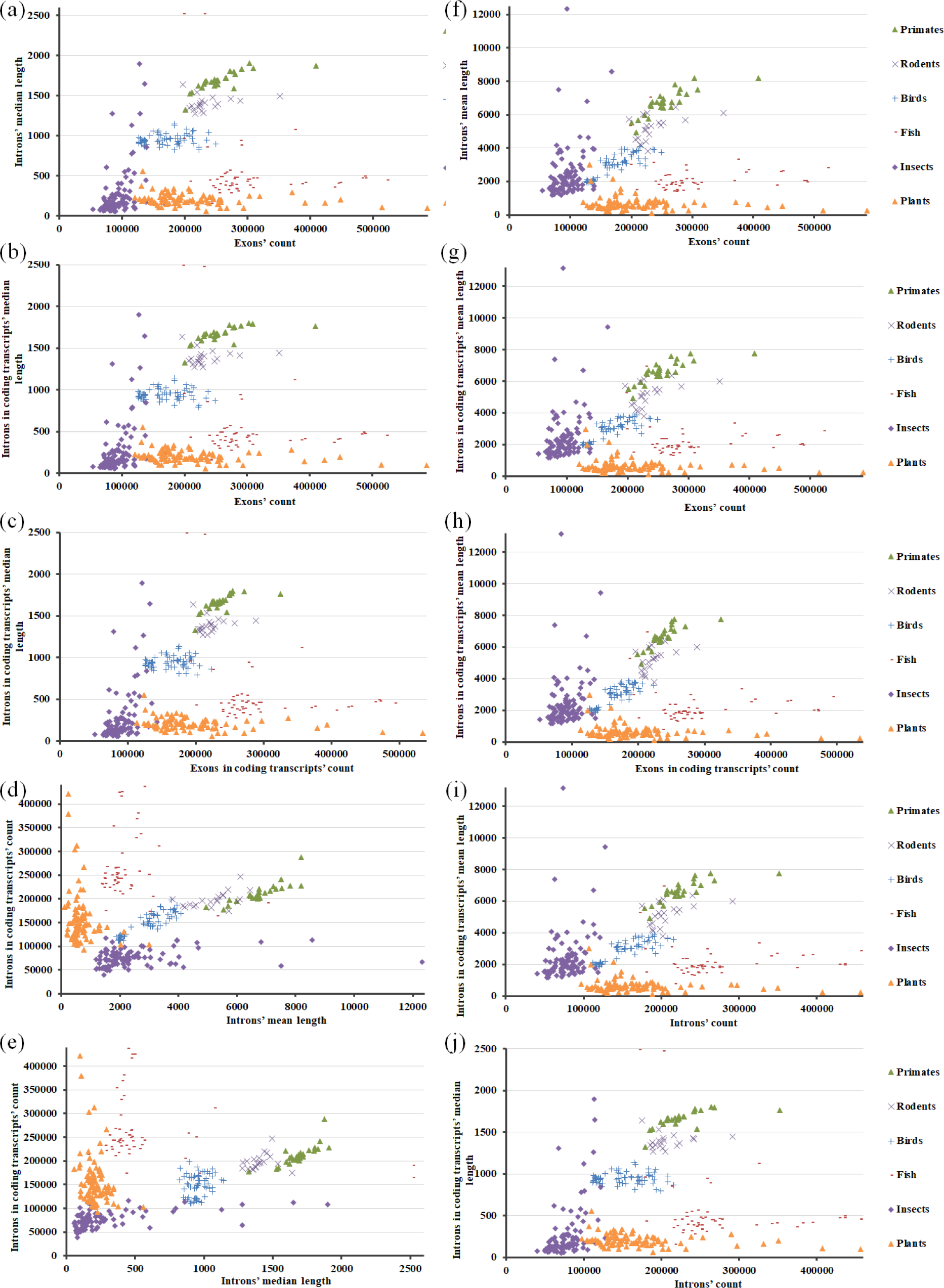
Scatter plots of the counts of exons *vs* median lengths of introns (a), counts of exons *vs* median lengths of introns in coding transcripts (b), counts of exons in coding transcripts *vs* median lengths of introns in coding transcripts (c), mean lengths of introns *vs* counts of introns in coding transcripts (d), median lengths of introns *vs* counts of introns in coding transcripts (e), counts of exons vs mean lengths of introns (f), counts of exons vs mean lengths of introns in coding transcripts (g), counts of exons in coding transcripts vs mean lengths of introns in coding transcripts (h), counts of introns vs mean lengths of introns in coding transcripts (i), and counts of introns vs median lengths of introns in coding transcripts (j), in eukaryotes.

Since no length-feature of exons is present in Fig. 7a,b,c,f,g,h with the exons’ data, corresponding plots with the mean or median values of exons are generated (Fig. 8). They reveal distinctive patterns (Fig. 8), but no separation of groups like in Fig. 7.

**Fig. 8.**
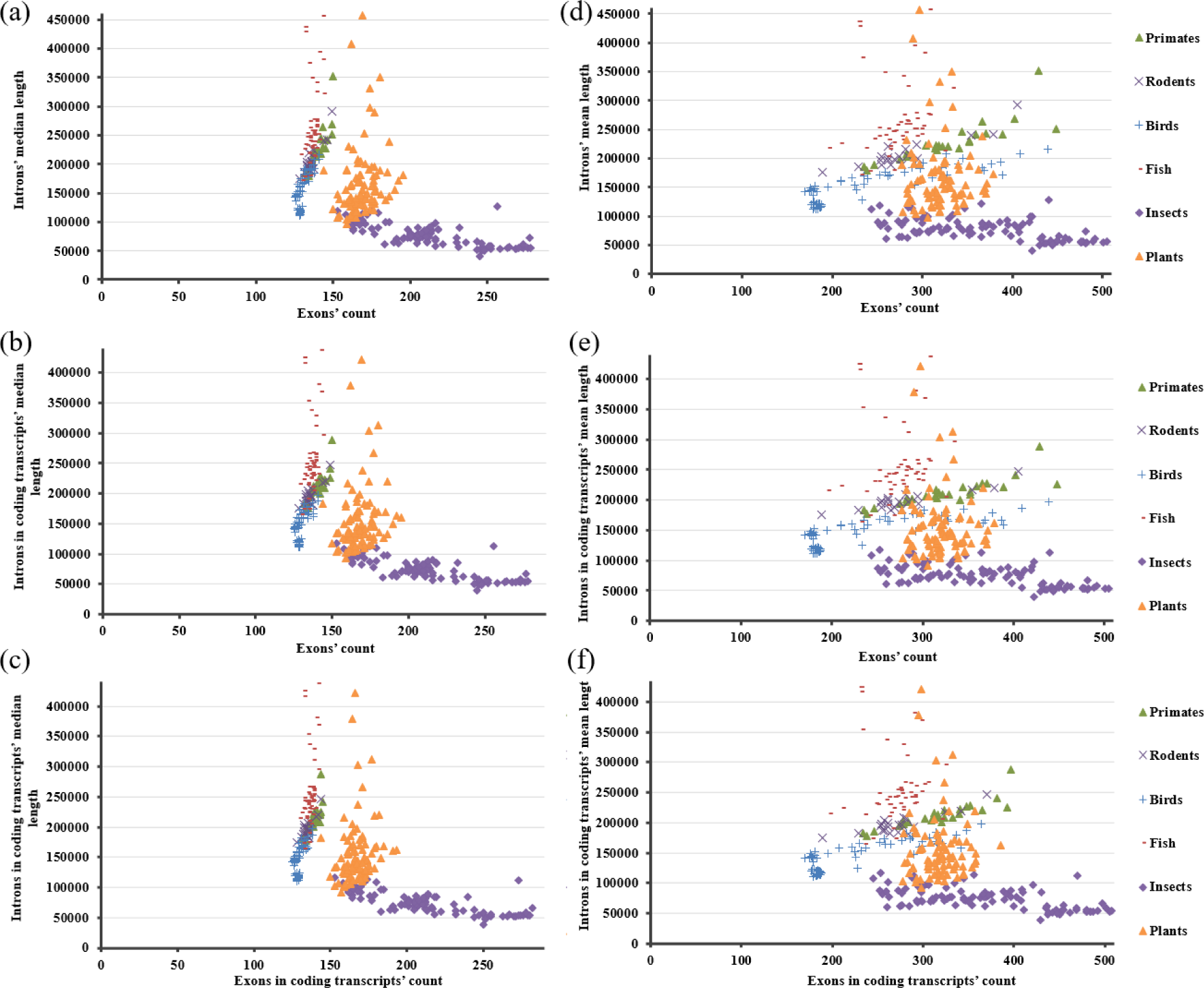
Scatter plots of the median lengths of exons *vs* counts of introns (a), median lengths of exons *vs* counts of introns in coding transcripts (b), median lengths of exons in coding transcripts *vs* counts of introns in coding transcripts (c), mean lengths of exons *vs* counts of introns (d), mean lengths of exons *vs* counts of introns in coding transcripts (e), mean lengths of exons in coding transcripts *vs* counts of introns in coding transcripts (f), in eukaryotes.

Missing group separation in Fig. 8 can be a standalone feature of the plotted data. Group-separation is observed also when the median lengths of introns or introns in coding transcripts are plotted against that of exons or exons in coding transcripts (Fig. 9).

**Fig. 9.**
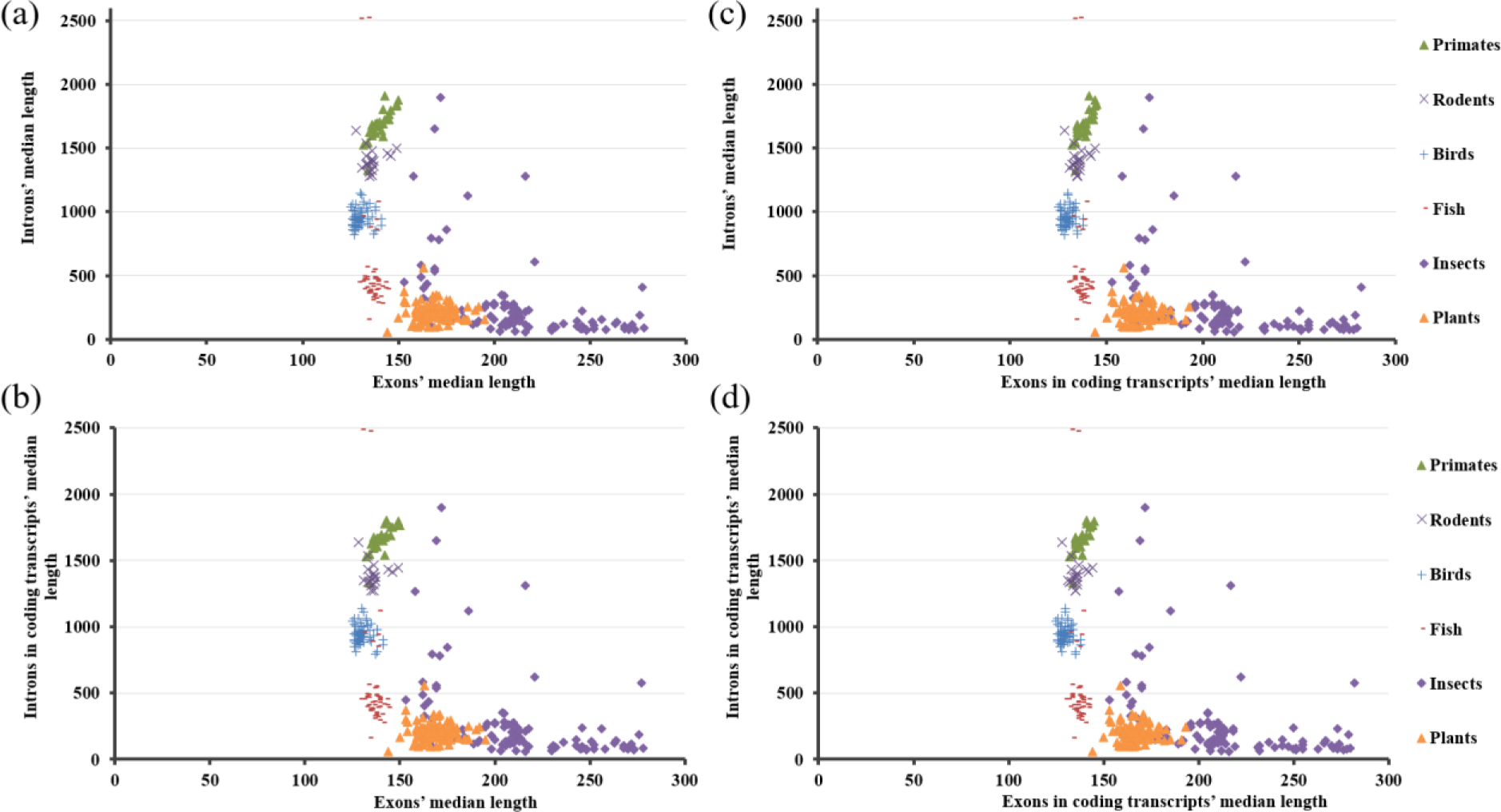
Scatter plots of the median lengths of exons *vs* introns (a), exons *vs* introns in coding transcripts (b), exons in coding transcripts *vs* introns (c), and exons in coding transcripts *vs* introns in coding transcripts (d), in eukaryotes.

### C. Pareto fronts in the cumulative data with max length values as the competing traits

Max lengths of the genes *vs* introns or introns in coding transcripts, and introns, or introns in coding transcripts *vs* introns in non-coding transcripts vary in a triangular constrained space (Fig. 10). This can well represent Pareto fronts of the traits under study (Sheftel, et al., 2013). We may consider only the plots with introns in coding transcripts and omit those with introns, as earlier (Fig. 10b,d), although the triangular shape is more well-defined in Fig. 10c. There are extremities, which can be the species maximizing both traits (Li, et al., 2019). However, it is of note that the extremities that are rather close to the borders are less in case of the plots with introns (Fig. 10a,c) than the plots with introns in coding transcripts (Fig. 10b,d). That observation is important here since we mentioned earlier that the plots of data with introns are like the plots of data with introns in coding transcripts. However, here we observe that the contribution of the introns in non-coding transcripts leads to a better Pareto front with less outliers (Fig. 10a,c compared to Fig. 10b,d). The borders generated by the Pareto front software (Fig. 11) include the points which are displayed as extremities in Fig.10. P-values with 10000 repetitions are ∼0.0004 and ∼0.0003, respectively, for the Pareto front analyses result with triangle fit depicted in Fig. 11a and Fig. 11b.

**Fig.10.**
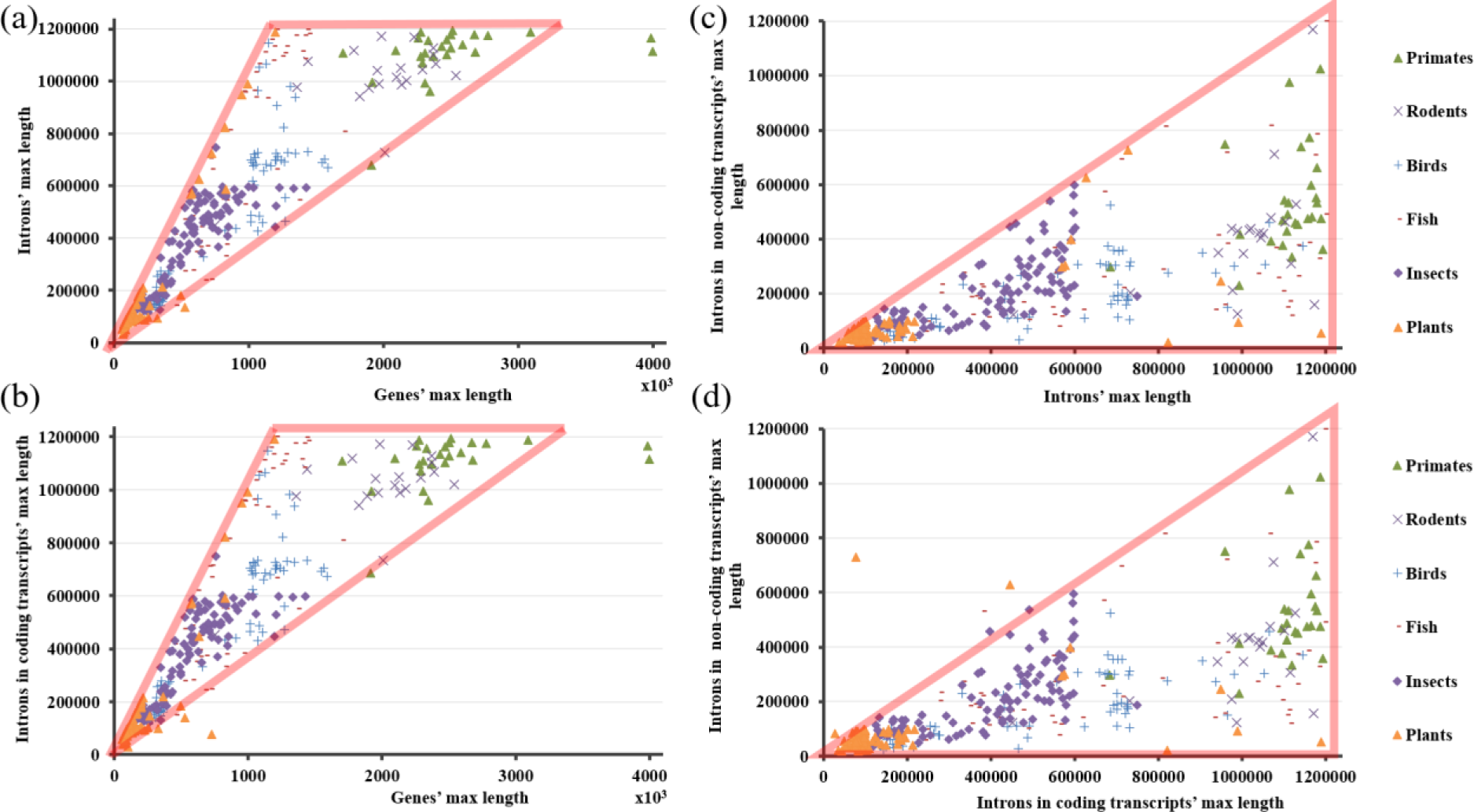
Scatter plots of the max lengths of genes *vs* introns (a), genes *vs* introns in coding transcripts (b), introns *vs* introns in non-coding transcripts (c), and introns in coding transcripts *vs* introns in non-coding transcripts (d), in eukaryotes. Triangular shapes are drawn onto the plotted data manually.

**Fig.11.**
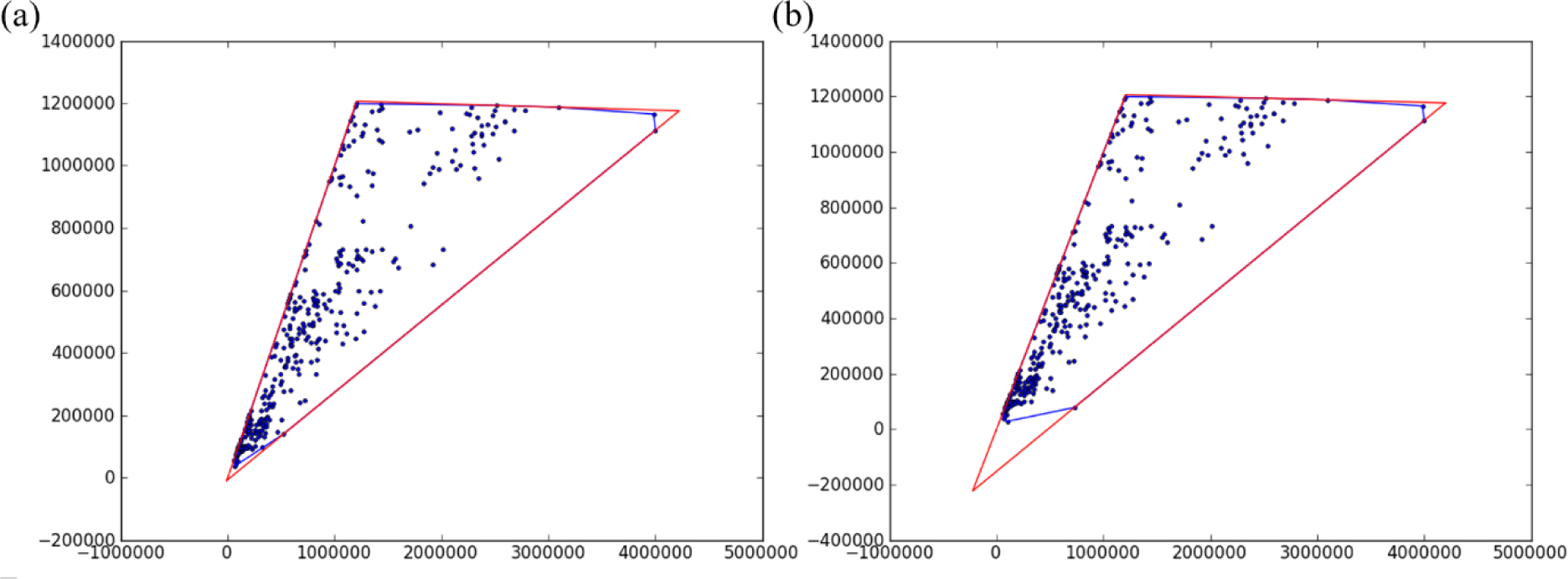
Pareto front analysis of the max lengths of genes *vs* introns (a) and genes *vs* introns in coding transcripts (c), in eukaryotes. As defined in the manual of the software, blue line indicates the convex hull of the data, and a red line indicates the best-fit triangle.

Other than the discussions above, one can interpret such triangular plot-shape as the constriction of data from being present outside of a hypothetical region with linear borders, which have positive, negative, or constant (independent) slopes. These hypothetical borders become limits of the parameters. In Fig. 10b, the triangularly shaped region has two positively correlated and one constant hypothetical-borders. Certain regions are populated by the data of distinct groups. Vertices represent specialists when the hypothetical edges are the pareto fronts of the competing traits (max lengths of genes, and introns in coding transcripts, or introns in coding, and introns in non-coding transcripts.) If we think in simple mathematical terms, this observation can also be interpreted as one of the parameters’ being constrained to be below, above, or equal to, the multiplier of another parameter, up to a certain value of the parameters if one or two of the edges is a constant line. As a result, we observed pareto fronts in the data with the max length values of the genes and introns in coding transcripts, as well as introns in coding and non-coding transcripts (Fig. 10). Same feature comparisons of the exons and exons in coding transcripts and of the exons and exons in coding transcripts are linear trait correlations (Supplemental_Fig_S1, respective plots at pages1–5). However, triangular space implies the involvement of three traits (Shoval, et al., 2012). Here, the third trait, which is not among the plotted parameters, but represented in the plot as the present relation, can be exons, mean lengths, but more likely the counts-data because counts of introns and exons contribute to the gene length. On the other hand, it can be inferred from Fig. 10b that the plotted-traits are not competing. Instead, they are positively correlated with different slopes, under the influence of the third trait. Species at the vertices are the specialists, which are optimized in one trait. Plants’, fishes’, and primates’ data are the vertices. Correlation of the max lengths of introns in coding transcripts is more positive towards the side of the fishes’, and less positive towards the side of the primates’ data. This relation can be seen through the numbers as well (Table 2) where the ratio of the max lengths’ range of the introns in coding transcripts and that of genes is 0.22 for primates. That ratio is not the highest for the fishes, it is the plants, but the genes’ max length is higher in the fishes than the plants (Table 2). Besides, data of plants is also going up along with the data of fishes at the left edge with the higher slope (Fig. 10a,b). The numbers in Table 2 explain the vertices and the approximate slopes, but the triangular shape is another related peculiarity.

**Table 2.** Max lengths’ ranges of the introns in coding transcripts and genes, their ratio, and the max lengths of genes and the introns in coding transcripts.

|  | **Introns in coding**<br>**transcripts' max**<br>**lengths' range (a)** | **Genes' max**<br>**lengths' range (b)** | **a/b** | **Genes' max**<br>**length** | **Introns in coding**<br>**transcripts' max**<br>**length** |
| --- | --- | --- | --- | --- | --- |
| **Primates** | 508833 | 2300158 | 0.22 | 4001895 | 1192712 |
| **Rodents** | 719773 | 1785683 | 0.40 | 2539466 | 1172235 |
| **Birds** | 1013993 | 1310512 | 0.77 | 1592281 | 1144717 |
| **Fishes** | 1051345 | 1257097 | 0.84 | 1704894 | 1198388 |
| **Insects** | 628839 | 1261140 | 0.50 | 1425452 | 748744 |
| **Plants** | 1161733 | 1136773 | 1.02 | 1196839 | 1189012 |

In conclusion, considering combined data, except that of plants and fishes in the counts-data, results of the current study are interpreted also as follows: Exons in the coding transcripts’ counts rather than their mean lengths increase with the mean lengths of genes while introns in the coding transcripts’ both mean lengths and counts, as well as the mean lengths of introns in the non-coding transcripts, increase with the genes’ mean lengths. This is also the case for the exons and introns, due to the dominance of exons in the coding transcripts and introns in the coding transcripts over those in the non-coding transcripts, respectively. The situation is similar in case of the median lengths. We also observed pareto fronts in the cumulative data with the max length values of genes and introns in coding transcripts, as well as introns in coding and non-coding transcripts.

## Materials and methods

Feature length information of the annotation reports individual species are from (Adiguzel, 2023). Namely, we collected the maximum (max) (*i*), minimum (min) (*ii*), median (*iii*), and mean (*iv*) lengths, and counts (*v*) of genes (*1*), all transcripts (*2*), mRNAs (*3*), misc_RNAs (*4*), tRNAs (*5*), lncRNAs (*6*), single-exon transcripts (*7*), coding transcripts (*8*), CDSs (*9*), exons (*10*), exons in coding transcripts (*11*), exons in non-coding transcripts (*12*), introns (*13*), introns in coding transcripts (*14*), and introns in non-coding transcripts (*15*). Missing of any feature length information was absent from this data, excluded during data collection. The number of primates is 26, rodents 20, birds 64, fishes 53, insects 99, and plants 90, making a total number of 352 eukaryotes. Focus of the current study is limited to the exons and introns. MS Excel is utilized for the plots, calculations, and statistics, unless otherwise stated. E.g., linear regression analysis of the data is performed with Graphpad Prism 8.2.1. (441). Normality of each distribution is calculated through the Jarque-Bera test, and the results are provided with the summary statistics, where p<0.05 was deemed as significant, i.e., not distributed normal (Supplemental_Table_S13). All plots’ document prepared with PPT productivity v3.1.0.13210.

As noted at the annotation reports’ pages, single-exon transcripts are the single-exon mRNAs, misc_RNAs, ncRNAs of class lncRNA; and excluding tRNAs, rRNAs, or ncRNAs of class other than lncRNA. Exons and introns have a similar explanation, wherein those in coding transcripts are mRNAs while those in non-coding transcripts are misc_RNAs and ncRNAs of class lncRNA. In relation to exons and introns, additionally it is stated that they are counted only once if shared by multiple transcripts. Again, in relation, the abbreviation NM stands for mRNA, XM stands for predicted mRNA, NR stands for non-coding RNA, and XR stands for predicted non-coding RNA. Mean length of genes is calculated as well, by assuming that the mean lengths of genes can also be approximated by taking the sum of the mean lengths of exons and introns, after multiplying them respectively with their counts per gene. Total length of intron per gene length is calculated by multiplying the mean length of introns with the average counts of introns per gene. Average counts of introns per gene are obtained by dividing the counts of introns to the respective counts of genes. Calculations are done with the same data.

Pareto front analysis is performed with the ParetoSW software (v 1.0) written by Hila Sheftel, Omer Ramote, and Oren Shoval, from Uri Alon Lab (www.weizmann.ac.il/mcb/UriAlon) (Grant, et al., 1985; Wilson, 1980; Norberg and Rayner, 1987). The analysis is performed as described in the user manual of the software, which is used for detection of polyhedron shape-matching variations that represent Pareto optimality of organisms regarding several evolutionary tasks, falling on low dimensionality polytopes like lines, triangles, etc.

## Acknowledgments

Ecology and evolutionary biology and biophysics societies of Turkey are acknowledged.

